# Crystal structure of the human mARC1 p.M187K variant

**DOI:** 10.64898/2026.05.26.727831

**Authors:** Charlotte M. Rehbein, Michel A. Struwe, Axel J. Scheidig

## Abstract

The human mitochondrial amidoxime reducing component 1 (mARC1) is a molybdenum-dependent enzyme whose protein-coding variants confer protection against common metabolic liver diseases. Whereas the frequent A165T variant acts largely through accelerated cellular degradation, the basis of protection by the rarer M187K variant remains obscure, as it has been reported that there are no differences between the “wild-type” and M187K variant protein in terms of cellular protein levels and localisation. Here, the crystal structure of the human mARC1 M187K variant, crystallised as a T4 lysozyme fusion after iterative micro-seeding, was determined at 1.63 Å resolution. The variant structure is essentially superimposable with the previously reported “wild-type” and A165T structures, with pairwise root-mean-square deviations of 0.3–0.4 Å, and the pentacoordinated molybdenum cofactor is fully intact. Differential scanning fluorimetry across a broad pH range revealed only a modest, pH-dependent decrease in thermal stability associated with the exchange, most pronounced at alkaline pH. These data suggest that the disease-protective effect of M187K is unlikely to originate from gross structural rearrangement or active-site perturbation.

**Synopsis:** The structure of the disease-relevant human mARC1 M187K variant was determined at 1.63 Å resolution after iterative micro-seeding. The variant does not display relevant perturbations of the overall protein fold or active site structure, but differential scanning fluorimetry detects a pH-dependent decrease in overall protein stability associated with the M187K amino acid exchange.

## 1. Introduction

The human mitochondrial amidoxime reducing component enzymes mARC1 and mARC2 (gene names *MTARC1, MTARC2*) are molybdoenzymes that were originally discovered due to their role in activating *N*-hydroxylated prodrugs of amidines, *e*.*g*., novel oral anticoagulants (Clement & Struwe, 2023, Havemeyer *et al*., 2006), and are recognised as a biotransformation pathway for pharmaceutical drugs (Klopp *et al*., 2024). Later, a potential involvement in lipid metabolism was suggested (Neve *et al*., 2012, Neve *et al*., 2015) and further supported by the phenotype observed in a murine knockout model, where mice with an *Mtarc2*^-/-^ genotype showed strong alterations in lipid metabolism compared to “wild-type” animals and a remarkable “resistance” towards high-fat diet-induced obesity (Rixen *et al*., 2019).

Interest in mARC enzymes has increased dramatically in recent years, after it was discovered that coding variants of mARC1 confer protection against common liver diseases like metabolic dysfunction-associated steatotic liver disease (MASLD) and metabolic dysfunction-associated steatohepatitis (MASH) (Emdin *et al*., 2020), including decreased liver-related mortality (Schneider *et al*., 2021). How mARC enzymes are linked to lipid metabolism and even what their physiological substrates are, and to which extent mARC1 and mARC2 are isofunctional, remain unknown (Struwe *et al*., 2023).

The most common variant detected among study populations is A165T and, consequently, this variant has received the most attention. mARC1 protein with the A165T amino-acid exchange has essentially the same structure as the “wild-type” protein (Struwe *et al*., 2022). However, in a biological context, mARC1 A165T is still considered a “loss of function” variant for practical purposes, as it shows increased cellular degradation of the protein and, thus, strongly depleted mARC1 protein levels (Dutta *et al*., 2024, Hou *et al*., 2024).

Genome-wide association studies have detected another single amino acid exchange variant, M187K, which is much rarer than A165T, but appears to confer protection against liver diseases in a very similar fashion. Interestingly, cell-culture models show that while *MTARC1* mRNA levels were similar for cells expressing “wild-type,” A165T- and M187K-variant mARC1, only A165T led to lower mARC1 protein levels and, potentially, mis-localisation of the protein. On the other hand, the M187K variant protein is detected at similar levels as the “wild-type” protein and is seen correctly co-localised with mitochondria (Wu *et al*., 2024, Smagris *et al*., 2024, Coyne *et al*., 2025).

Early *in vitro* characterisation of heterologously expressed mARC variants did not identify A165T or M187K as protein variants with markedly impaired molybdenum-cofactor loading or *N*-reductive activity against the model substrate benzamidoxime (Ott *et al*., 2014). For the A165T protein variant, it is by now well-established that accelerated protein degradation leads to lower effective protein levels *in cellulo*, and this is thought to be central to the lipid phenotype. The picture is much less clear for the rarer M187K variant that displays a similar phenotype, but no changes in degradation or localisation.

Here, we determined the crystal structure of the mARC1 M187K protein variant at near-atomic resolution. While obtaining diffraction-quality crystals was somewhat more challenging than for the “wild-type” and A165T variants, the structure itself is virtually indistinguishable from previously reported mARC1 structures.

## 2. Materials and methods

Standard methods were used for all molecular biology procedures (Ausubel *et al*., 2003). Unless stated otherwise, all chemicals were purchased from Carl Roth GmbH+Co. KG (Karlsruhe, Germany) and were of > 99 % purity, or of the highest purity available.

### 2.1. Macromolecule production

The pQE80-based expression plasmid for the hmARC1-T4L fusion protein carrying the p.M187K variant was generated by site-directed mutagenesis of the previously described plasmid (Kubitza, Ginsel, *et al*., 2018). Mutagenesis was achieved using the Q5 site-directed mutagenesis kit (New England Biolabs #E0554) according to the manufacturer’s instructions with the mutagenic primers 5’-CGAGCCTCACA<u>AA</u>CGACCGAGAC-3’ and 5’-AAGTGCACCAGGCGG-3’, which had been designed using the *NEBaseChanger* web tool (https://nebasechanger.neb.com). Successful mutagenesis was confirmed by Sanger sequencing using the primers 5’-GTATCACGAGGCCCTTTCGTCT-3’ and 5’-CATTACTGGATCTATCAACAGGAG-3’.

Heterologous expression was performed in the molybdopterin-accumulating *Escherichia coli* strain TP1001 (Palmer *et al*., 1996). Briefly, bacteria were grown at 37 °C in terrific broth supplemented with 1 mM Na_2_MoO_4_, 0.1 mM ferric citrate, 0.5 mM MgSO_4_, 0.5 mM MgCl_2_, 5 mM NH_4_Cl and 100 µg × ml^-1^ carbenicillin until an apparent OD600 of ≈ 0.2 was measured. The incubation temperature was reduced to 22 °C and 0.05 mM isopropyl β-D-thiogalactopyranoside was added to induce expression. After 22 hours, cells were harvested by centrifugation, flash-frozen in liquid nitrogen and stored at – 80 °C until further use.

Cells were resuspended in purification buffer (50 mM NaH_2_PO_4_, 300 mM NaCl, 10 mM Na_2_MoO_4_, 10 % w/v glycerol, pH 7.4) supplemented with 0.7 mM phenylmethylsulfonyl fluoride (PMSF) and a catalytic amount of deoxyribonuclease I (VWR #A3778). Cells were lysed by two passages through an Emulsiflex C3 high-pressure homogenizer (Avestin) and cell debris was removed by centrifugation. The lysate was loaded onto a HiTrap TALON crude 1 mL column (Cytiva # 29048565) equilibrated in purification buffer and connected to an ÄKTA pure 25 M low-pressure chromatography system (Cytiva). Non-specifically bound proteins were washed from the column with 10 mL of 15 mM imidazole in purification buffer, followed by elution of the target protein with a linear gradient from 15 – 375 mM imidazole over 20 mL. Fractions were analysed by SDS-PAGE, pooled, concentrated using a spin concentrator (Carl Roth #25TL) and exchanged into storage buffer (50 mM TRIS-HCl, 150 mM NaCl, 5 % w/v glycerol, pH 6.8) using a PD-10 desalting column (Cytiva #17085101).

### 2.2. Crystallisation

In contrast to the T4 lysozyme-fusion proteins of “wild-type” mARC1 and its A165T variant, the M187K variant protein did not crystallise spontaneously. Diffraction-quality crystals were obtained through cross-seeding with the “wild-type” construct. All crystals were obtained by hanging-drop vapour diffusion on glass cover slips (Carl Roth #P235.1) over 24-well crystallisation plates (Molecular Dimensions #MD3-11), sealed with silicone grease (#Carl Roth #0856.1). The “wild-type” protein was crystallised against a reservoir containing 100 mM bis-tris propane-HCl pH 6.5, 20 mM sodium molybdate, 27.5 % (w/v) PEG 3350. The crystal suspension was crushed using a rounded tip glass rod and transferred into a 1.5 mL reaction tube containing 100 µL freshly prepared reservoir to create a seed stock, and a serial dilution in 1:10 steps was prepared. These seed stocks were used to initiate crystallisation of M187K-variant mARC1-T4L in hanging drops containing 2 µL hmARC1 protein solution (c = 6 mg × mL^-1^), 1 µL reservoir and 1 µL seed solution. Crystals obtained from these droplets were once again used to prepare seed stocks, using an optimised reservoir composition: 100 mM bis-tris propane-HCl pH 8.0, 20 mM Na_2_MoO_4_, 27.5 % PEG3350, 10 mM tris-(2-carboxyethyl)-phosphine hydrochloride (TCEP), and crystals were obtained from droplets prepared from 2.5 µl protein solution, 1 µl of reservoir solution and 0.5 µl of seed solution. These crystals were subsequently used for another round of seeding to obtain crystals suitable for X-ray diffraction experiments.

### 2.3. Data collection and processing

An intergrown crystal (see figure 1C) was broken, and one fragment was transferred into a 5 µl droplet of fresh reservoir solution. After approx. 45 s, the crystal was mounted into a CryoLoop (Hampton Research) and flash-cooled in liquid nitrogen. X-ray diffraction data were collected at the P13 beamline operated by EMBL at DESY, Hamburg (Cianci *et al*., 2017), equipped with an EIGER 16M detector (Dectris). Diffraction data were indexed and integrated with *XDS* (Kabsch, 2010), followed by scaling and merging with *AIMLESS* (Evans & Murshudov, 2013). The high-resolution limit was set to 1.63 Å, where the outer-shell ⟨I/σ(I)⟩ is 2.2, and CC_1/2_ is 0.790.

**Figure 1.**
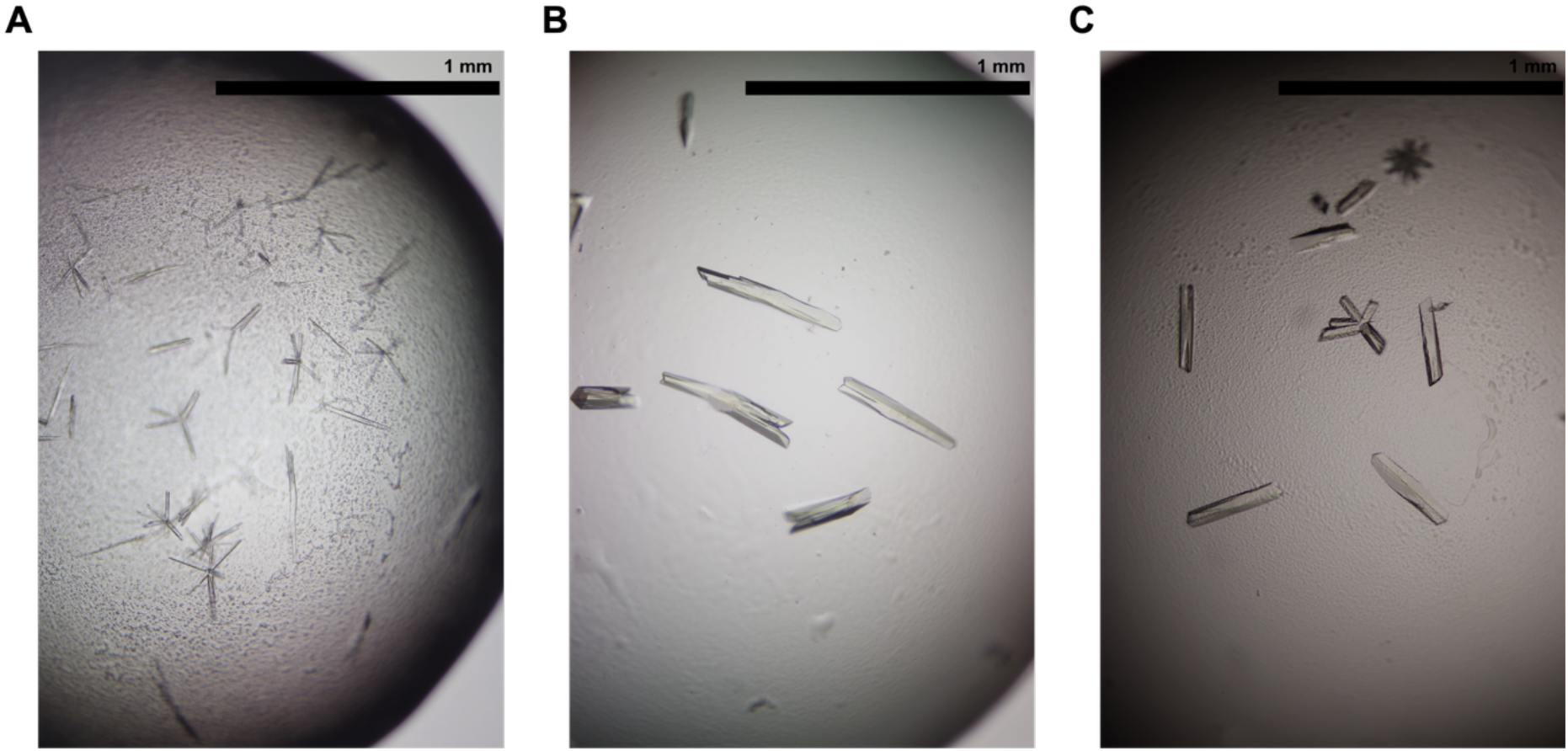
Crystals formed by M187K-variant mARC1-T4L. (A) After micro-seeding with crushed crystals of “wild-type”-mARC1-T4L, (B) after micro-seeding with crushed crystals from the droplet shown in Panel A and, (C) after micro-seeding with crushed crystals from the droplet shown in panel B. Crystal morphology gradually improves over multiple rounds of seeding. The crystal used for structure determination was a fragment of the intergrown crystal seen at the centre of panel C.

### 2.4. Structure solution and refinement

Phasing was achieved by molecular replacement with *Phaser* (McCoy *et al*., 2007) using the structure of “wild-type” mARC1-T4L (pdb_00006fw2). The structure was refined using *PHENIX AutoBuild* (Terwilliger *et al*., 2008), *phenix*.*refine* (Afonine *et al*., 2012) and manual manipulations in *COOT* (Emsley *et al*., 2010). The model and structure factors have been deposited in the RCSB Protein Data Bank under the accession code pdb_000029aj.

### 2.5. Differential scanning fluorimetry

In order to assess the impact of the M187K protein variant on the overall stability of mARC1, we compared “wild-type” and M187K-variant proteins by differential scanning fluorimetry (DSF). For these assays, mARC1 constructs without the internal T4 lysozyme fusion were used. Site-directed mutagenesis was achieved using the same primers as for the crystallisation construct and protein was purified by immobilised metal affinity chromatography as described previously. For DSF analysis, the protein was diluted to a concentration of approx. 1 mg × mL^-1^ in storage buffer. 4 µl of 5,000 × SYPRO Orange (Thermo Fisher #S6650) were added to 1.1 mL of protein solution (final concentration ca. 18 ×). 10 µl of this solution were distributed into a 96-well RT-PCR plate (Sarstedt #72.1979.102) and mixed with 10 µl buffer concentrates from the Durham pH Screen (Molecular Dimensions #MD1-101), before sealing the plate with a transparent film (Sarstedt 95.1999). Melting curves were recorded through the TAMRA channel of an ABI7300 RT-PCR instrument (Thermo Fisher). Briefly, fluorescence intensity was measured between 21 °C and 75 °C, with 0.25 min equilibration time at each temperature followed by 1 min readout time. Melting temperatures were determined as the inflection point of a Boltzmann sigmoidal curve fitted to the data using *Excel for Mac* 16.107 (Microsoft) and *Prism for macOS* 11.0.0 (GraphPad), following a standard procedure described elsewhere (Huynh & Partch, 2015). Assays were performed as technical duplicates to assess reproducibility of results.

## 3. Results and discussion

### 3.1 Diffraction-quality crystals of M187K-variant mARC1 protein are obtained after multiple rounds of micro-seeding

In contrast to “wild-type” (Kubitza, Ginsel, *et al*., 2018) and A165T (Struwe *et al*., 2022) mARC1-T4L fusion proteins, our preparation of the M187K variant of the same construct did not initially generate any crystals in hanging-droplet vapour diffusion experiments with solutions that had worked for the other two variants. However, the M187K variant did form crystals when a seed stock was prepared from crystals of the “wild-type” construct. Initially, these cross-seeded crystals were very thin needles with visible growth defects (figure 1A). Size and morphology improved markedly when these crystals were used for another round of seeding (figure 1B) and diffraction-quality crystals were eventually obtained from a third round of seeding (figure 1C).

### 3.2 “Wild-type”, M187K- and A165T-variant proteins share the same three-dimensional structure

The structure of the M187K-variant of mARC1 was solved by molecular replacement and refined, with refinement statistics shown in table 4.

**Table 1.**
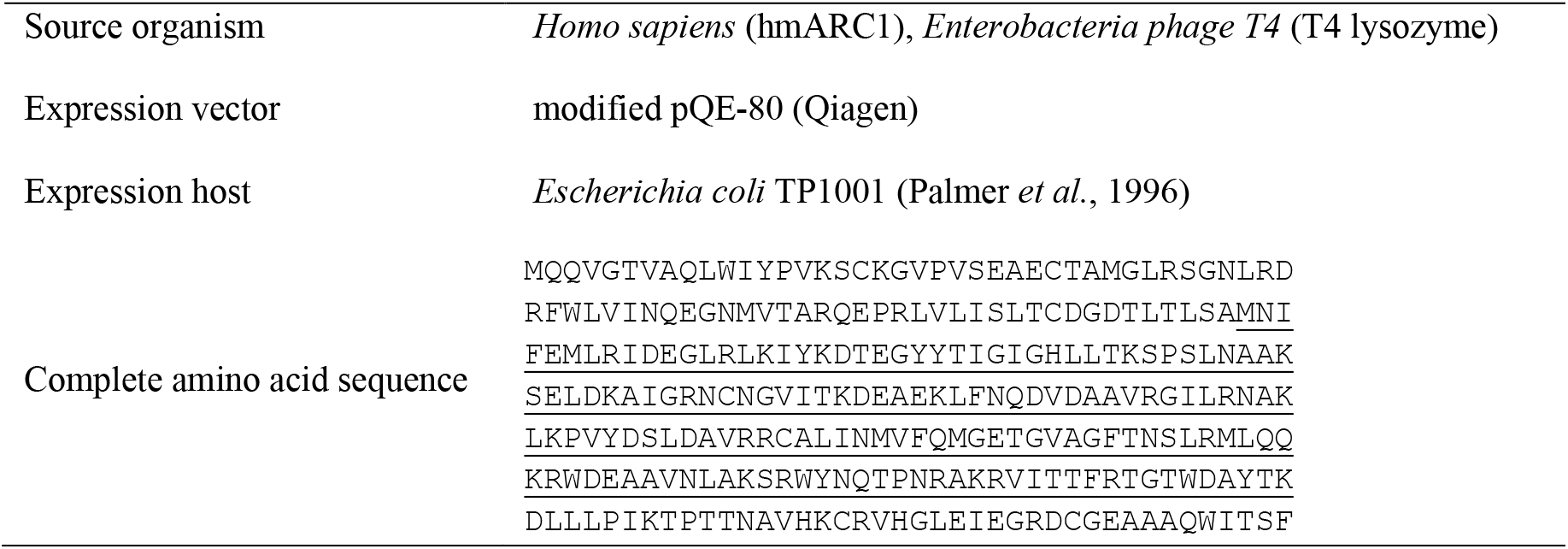
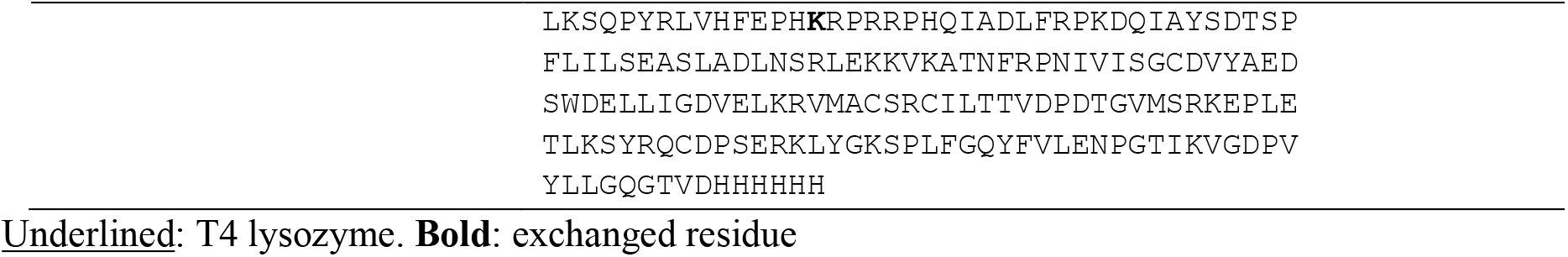
Macromolecule production information.

**Table 2.**
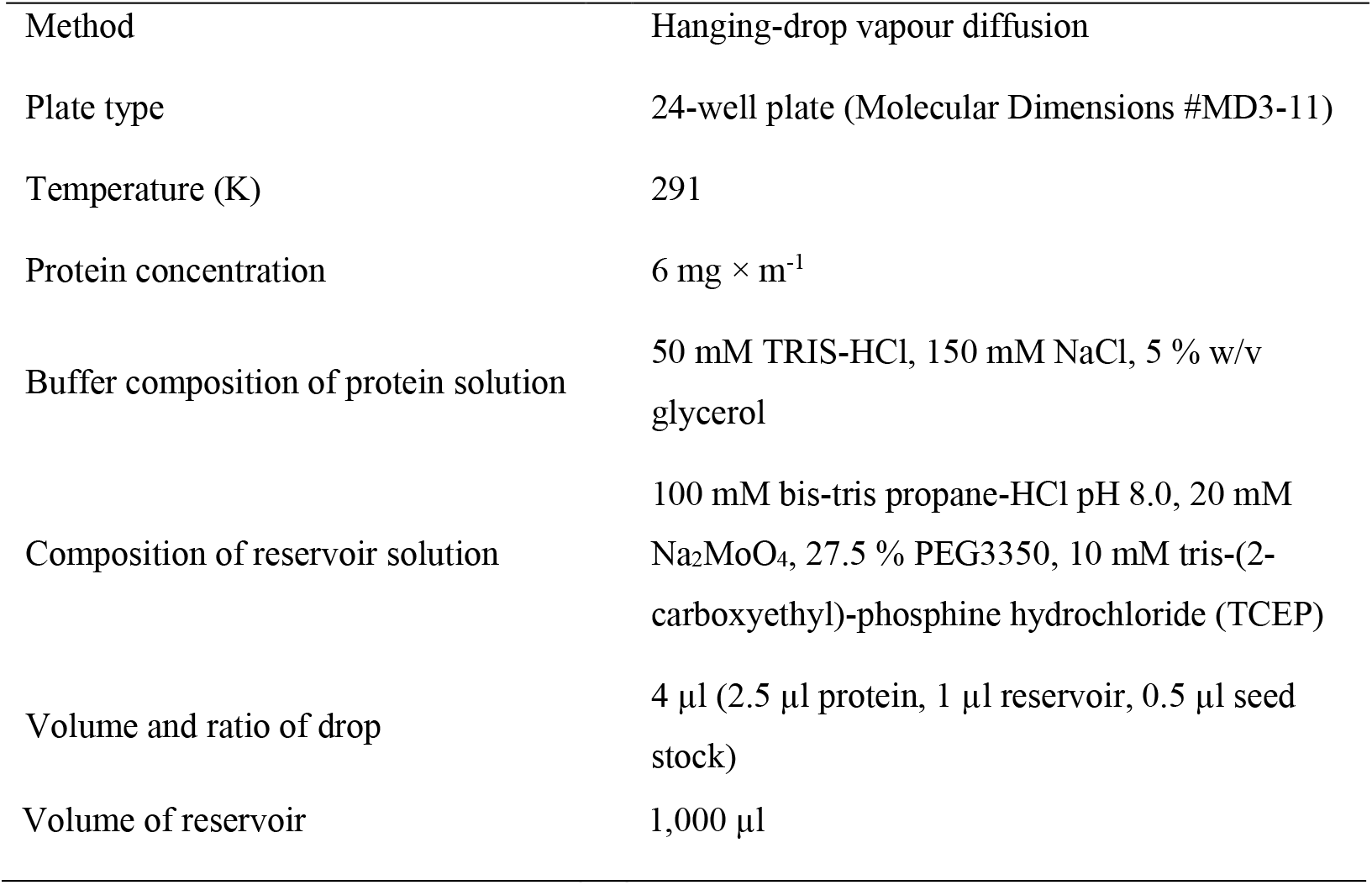
Crystallisation.

|  |  |
| --- | --- |
| Method | Hanging-drop vapour diffusion |
| Plate type | 24-well plate (Molecular Dimensions #MD3-11) |
| Temperature (K) | 291 |
| Protein concentration | $6 \text{ mg} \times \text{m}^{-1}$ |
| Buffer composition of protein solution | 50 mM TRIS-HCl, 150 mM NaCl, 5 % w/v glycerol |
| Composition of reservoir solution | 100 mM bis-tris propane-HCl pH 8.0, 20 mM Na<sub>2</sub>MoO<sub>4</sub>, 27.5 % PEG3350, 10 mM tris-(2-carboxyethyl)-phosphine hydrochloride (TCEP) |
| Volume and ratio of drop | 4 µl (2.5 µl protein, 1 µl reservoir, 0.5 µl seed stock) |
| Volume of reservoir | 1,000 μl |

**Table 3.** Data collection and processing. Values for the outer shell are given in parentheses.

|  |  |
| --- | --- |
| Diffraction source | P13 (PETRA III DESY) |
| Wavelength (Å) | 0.9762 |
| Temperature (K) | 100 |
| Detector | EIGER X 16 M detector (DECTRIS) |
| Crystal-detector distance (mm) | 229.328 |
| Rotation range per image (°) | 0.1 |
| Total rotation range (°) | 360 |
| Exposure time per image (s) | 0.01 |
| Space group | $P2_12_12_1$ |
| $a, b, c$ (Å) | 60.99, 75.07, 110.66 |
| $\alpha, \beta, \gamma$ (°) | 90, 90, 90 |
| Mosaicity (°) | 0.121 |
| Resolution range (Å) | 62.12–1.63 |
| No. of unique reflections | 64,043 |
| Completeness (%) | 99.9 (99.7) |
| Redundancy | 13.6 |
| $\langle I/\sigma(I) \rangle$ | 8.5 (2.2) |
| $R_{merge}$ | 0.190 (1.148) |
| $CC(1/2)$ | 0.989 (0.790) |
| Overall $B$ factor from Wilson plot ($\text{\AA}^2$) | 25.56 |

**Table 4.** Structure solution and refinement.

|  |  |
| --- | --- |
| Resolution range ($\text{\AA}$) | 62.12–1.63 (1.66–1.63) |
| Completeness (%) | 99.9 (99.7) |
| No. of reflections, working set | 60912 (2714) |
| No. of reflections, test set | 3131 (136) |
| Final $R_{\text{cryst}}$ | 0.188 (0.239) |
| Final $R_{\text{free}}$ | 0.215 (0.283) |
| No. of non-H atoms | 4214 |
| Protein | 3563 |
| Ligand | 67 |
| Water | 584 |
| R.m.s. deviations |  |
| Bonds ($\text{\AA}$) | 0.010 |
| Angles ($^\circ$) | 1.054 |
| Average $B$ factors ($\text{\AA}^2$) | |
| Protein | 30.5 |
| Ligand | 35.5 |
| solvent | 37.1 |
| Ramachandran plot |  |
| Most favoured (%) | 97.5 |
| Allowed (%) | 2.5 |
| Outliers (%) | 0.00 |

We subsequently performed structural comparisons of the three mARC1 variants using the *DALI* webserver (Holm *et al*., 2023). The RMSD values for the pairwise comparison of the M187K-variant (*this study*) with “wild-type” mARC1 (PDB: pdb_00006fw2) and A165T-variant mARC1 (PDB: pdb_00007p41) are reported at 0.4 Å and 0.3 Å, respectively. Visual inspection of the overlaid models shows that, apart from the side chain of M187 / K187, the three structures are virtually identical to each other (figure 2A). In order to verify that the crystallised species is the M187K variant, both lysine and methionine were modelled and refined for residue 187. When methionine (“wild-type”) rather than lysine (variant of interest) is fitted, a strong negative peak appears in the difference map, confirming that we are indeed looking at crystals of the M187K variant protein (figure 2B, C). Electron density for the prosthetic group shows an intact pentacoordinated [MoO_2_(S_Cys_)(PDT)]^−^ cofactor, which is consistent with structures derived from spectroscopic data on human mARC1 and its *E. coli* homologue YcbX (Struwe *et al*., 2024, Yang *et al*., 2023) and previously published X-ray crystal structures of mARC1 variants (Struwe *et al*., 2022, Kubitza, Bittner, *et al*., 2018). Overall, the active site of mARC1 seems unaffected by the M187K amino acid exchange (figure 2D). Given that the distance between the catalytic Mo ion and the Cα atom of M187/K187 is some 26 Å (similar to the distance between the Mo ion and A165/T165), it is not necessarily expected that the amino acid substitution would affect the active site. Nonetheless, it is worth documenting that no such perturbation is observed.

**Figure 2.**
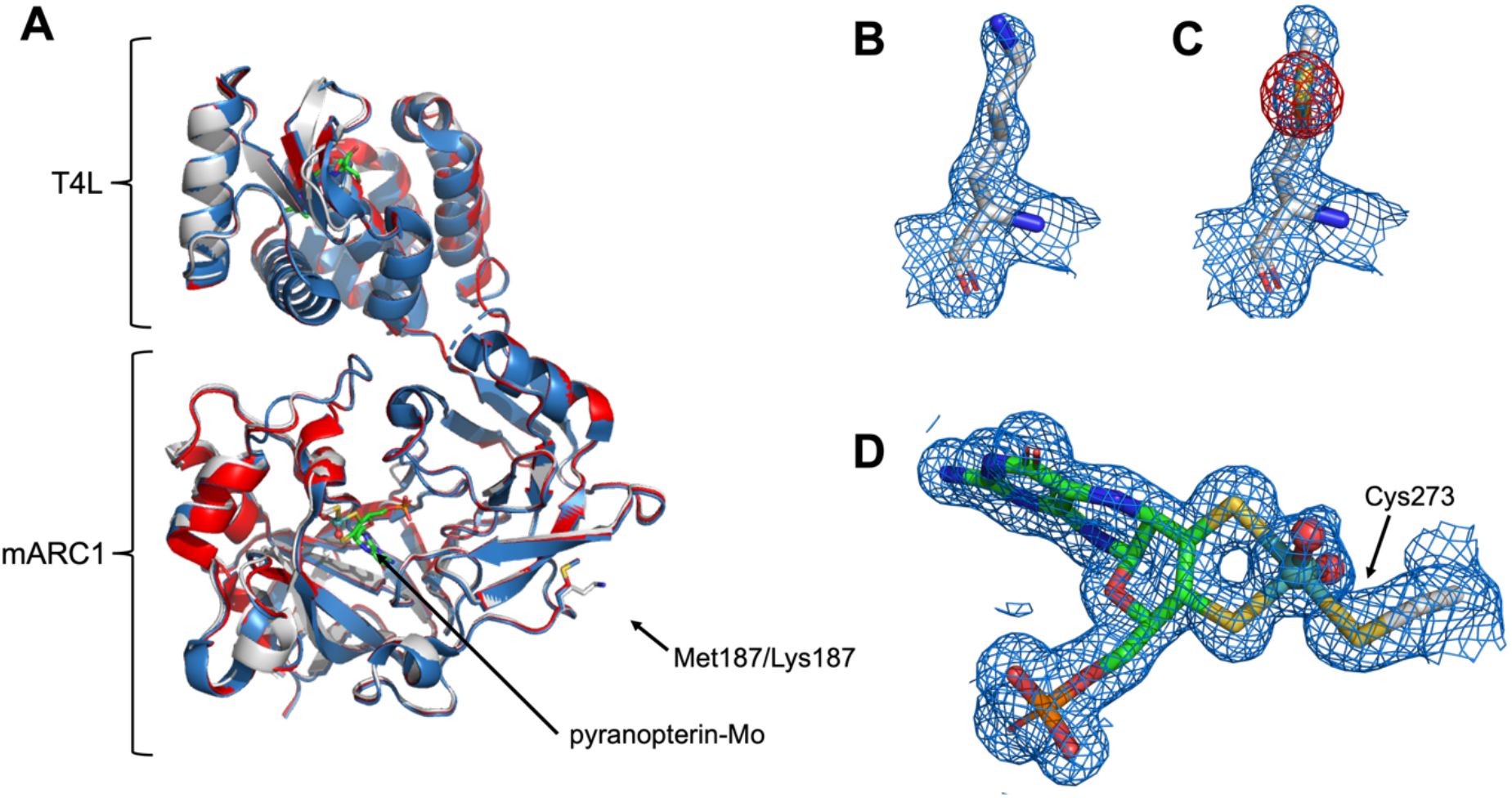
(A) Overlay of the structure reported here (grey, M187K variant), pdb_00006fw2 (blue, mARC1 “wild-type”) and pdb00007p41 (red, mARC1 A165T variant). The structures are virtually indistinguishable from one another with the only relevant differences being observed at the actual mutation sites. (B/C) Electron density maps for the side chain of residue 187, when (B) the residue is a lysine during refinement or (C) the residue is refined as a methionine (according to the “wild-type” species). 2Fo-Fc maps are displayed as a blue mesh and a contour level of 1σ. Fo-Fc maps are displayed as red (negative density) and green (positive density) meshes at a contour level of 4σ. If this residue is refined as a methionine, a strong negative peak at the methionine thioether sulfur (S^δ^) is observed. (D) 2Fo-Fc electron density map in which the pyranopterin-Mo cofactor has been fitted shows the prosthetic group fully intact.

### 3.3 The M187K amino acid exchange has a modest pH-dependent effect on thermal stability

The Durham pH screening kit comprises 93 different buffer conditions (28 individual buffer substances, spanning a pH range from 4 – 11). mARC1 displays optimal stability at acidic or near-neutral pH, with melting temperatures decreasing noticeably at alkaline pH. This general stability profile is observed both for the “wild-type” and M187K-variant proteins. However, we do notice that the M187K variant appears generally *less* stable compared to the “wild-type” protein. The effect is small at acidic pH but becomes stronger as pH increases. Example melting curves at different pH values are presented in figure 3. All estimated melting temperatures and their correlation with buffer pH are displayed in supplementary figure S1/S2.

**Figure 3.**
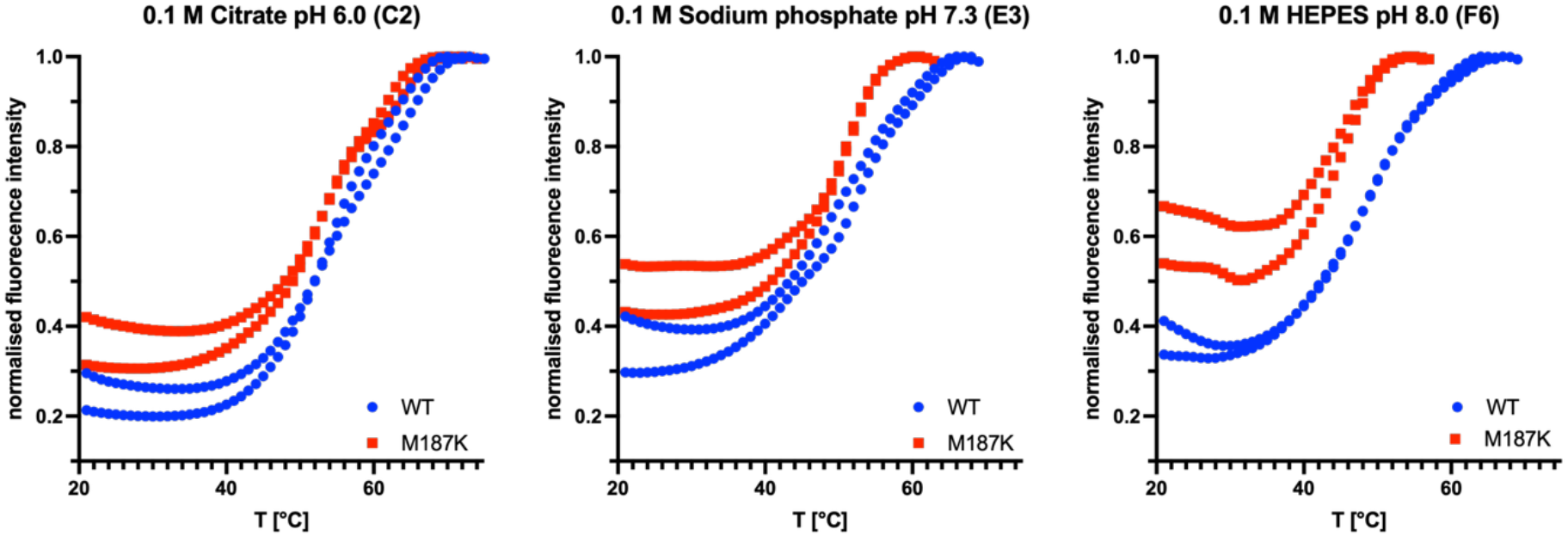
Example melting curves of “wild-type” (WT) and M187K-variant mARC1 at three example pH values. Blue points represent melting curves recorded for the “wild-type” protein and red squares represent melting curves recorded for the M187K variant protein. For clarity, curves were normalised to the peak intensity. Curves recorded for the M187K variant are shifted to lower temperatures and generally display higher initial fluorescence, consistent with the amino acid exchange decreasing thermal stability. This effect becomes more pronounced as pH increases.

While these data suggest a modest destabilising effect of the M187K amino acid exchange, it should be noted that at pH values likely to be biologically relevant (6 – 8) both “wild-type” and M187K-variant mARC1 can be considered to be quite *stable* proteins, with estimated melting temperatures of the protein generally higher than 45 °C. When *N*-reductive activities of various mARC1 protein variants, including “wild-type” and M187K, were investigated using *in vitro* enzyme activity assays, these were performed at optimised reaction conditions in 20 mM 2-(*N*-morpholino)ethanesulfonic acid (MES), pH 6.0, at 37 °C, and no significant differences in enzymatic activity (determined as generation of HPLC-quantified benzamidine from benzamidoxime) were observed (Ott *et al*., 2014). This is consistent with observations made by DSF, *i*.*e*., both variants being very stable in acidic MES buffer (T_m_ > 50 °C).

In summary, DSF data do not support the conclusion that decreased thermal stability of the M187K variant would be responsible for a loss of function under physiological circumstances that would clarify the variant’s influence on hepatic lipid accumulation. The slight decrease in thermal stability may, however, explain why multiple rounds of micro-seeding were necessary in order to obtain crystals that diffracted to high resolution for the M187K variant, consistent with reports that DSF can be predictive for crystallisation behaviour and diffraction quality (Sato & Senda, 2026).

## Conclusion

In conclusion, the M187K substitution does not induce a detectable rearrangement of the mARC1 fold or molybdenum cofactor site in the crystallographic construct used here. The variant does, however, display altered crystallisation behaviour and a modest pH-dependent reduction in thermal stability. The presented data do not support the hypothesis that the disease-protective phenotype of the M187K amino acid exchange is caused by large-scale structural disruptions or perturbations of the active site.

## Acknowledgements

We gratefully acknowledge access to the core facilities of the BiMo/LMB of Kiel University and thank Brigitte Bittner for technical assistance. The synchrotron data were collected at the P13 beamline operated by EMBL Hamburg at the PETRA III storage ring (DESY, Hamburg, Germany). We would like to thank Michael Agthe for the assistance in using the beamline.

GPT-5.5 was used for assistance with spelling, grammar and stylistic refinement. All scientific content, interpretation and conclusions are the responsibility of the authors.

## Supporting information

**Supplementary figure S1.**
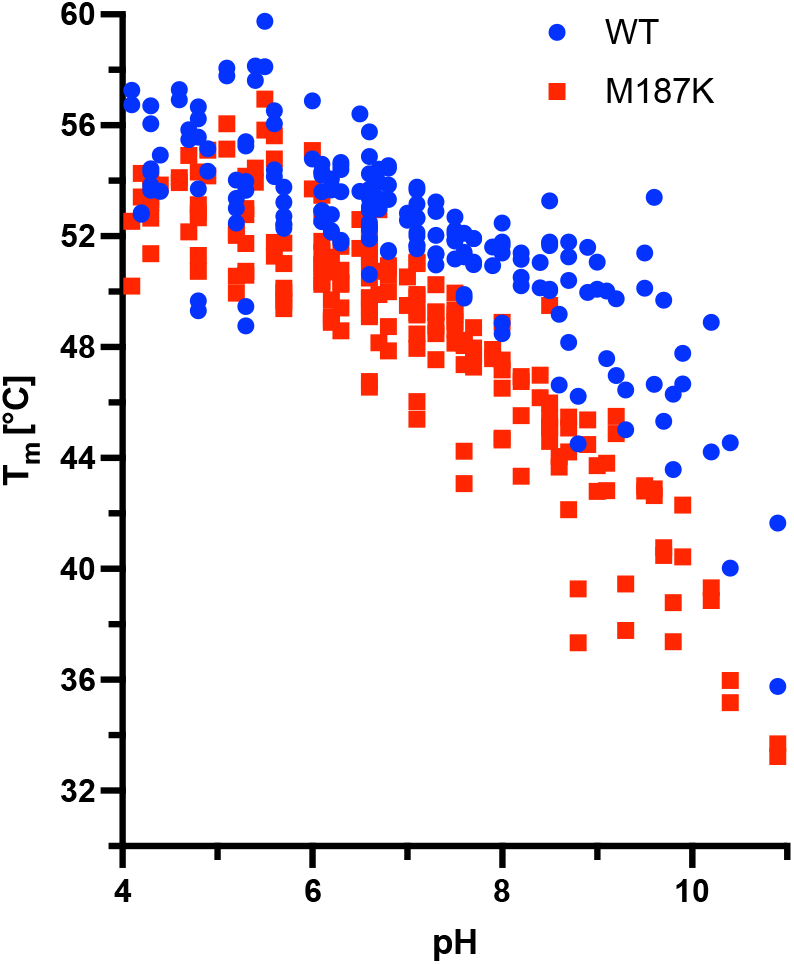
Melting temperatures of “wild-type” and M187K-variant mARC1 (estimated by Boltzmann-fitting of the melting curves) plotted against the respective buffer pH. Replicate values are displayed as separate points.

**Supplementary figure S2.**
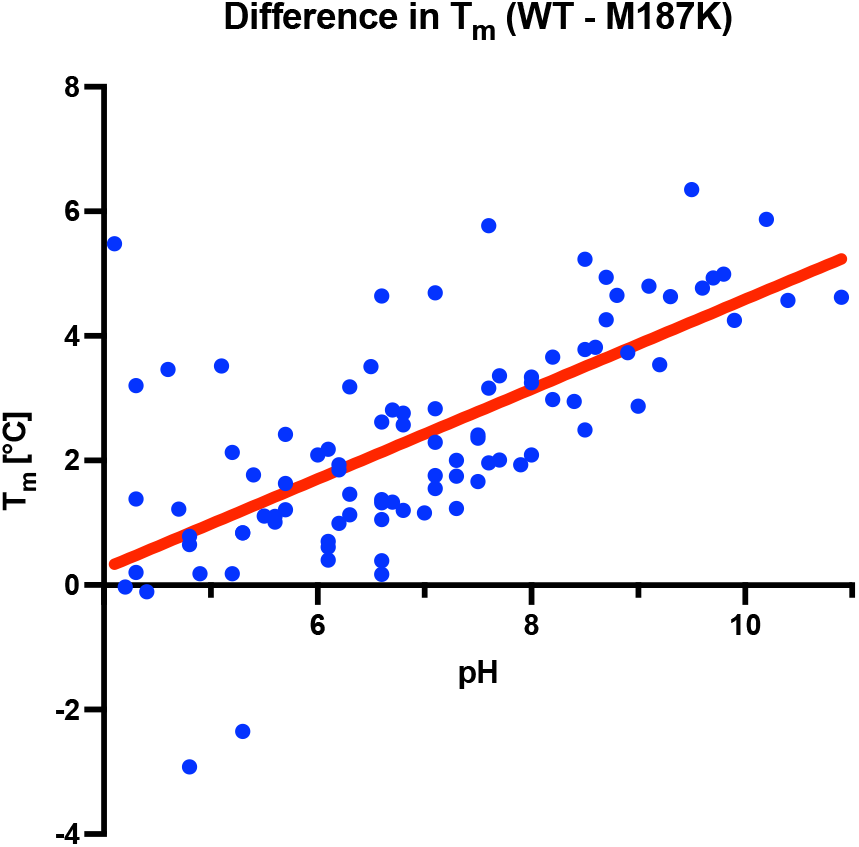
Differences in fitted T_m_ (average of two replicates; “wild-type” minus M187K variant) displayed as blue dots. A trend line (red) shows that with very few outliers, the differences in T_m_ tend to increase with pH. The trend line has an r^2^ of 0.566 and the P value for the slope being non-zero is reported as 0.0001 by *Prism for macOS* 11.0.0 (GraphPad).

